# Neo-sex chromosomes in the Monarch butterfly, *Danaus plexippus*

**DOI:** 10.1101/036483

**Authors:** Andrew J. Mongue, Petr Nguyen, Anna Volenikova, James R. Walters

**Affiliations:** Department of Ecology and Evolutionary Biology, University of Kansas, Lawrence, KS, USA; Faculty of Science, University of South Bohemia, 370 05 České Budějovice, Czech Republic; Institute of Entomology, Biology Centre CAS, 370 05 České Budějovice, Czech Republic

**Keywords:** sex chromosomes, evolution, Lepidoptera, genomics, chromosomal fusion

## Abstract

We report the discovery of a neo-sex chromosome in Monarch butterfly, *Danaus plexippus*, and several of its close relatives. Z-linked scaffolds in the *D. plexippus* genome assembly were identified via sex-specific differences in Illumina sequencing coverage. Additionally, a majority of the *D. plexippus* genome assembly was assigned to chromosomes based on counts of 1-to-1 orthologs relative to the butterfly *Melitaea cinxia* (with replication using two other lepidopteran species), in which genome scaffolds have been mapped to linkage groups. Sequencing-coverage based assessments of Z-linkage combined with homology based chromosomal assignments provided strong evidence for a Z-autosome fusion in the *Danaus* lineage, involving the autosome homologous to chromosome 21 in *M. cinxia*. Coverage analysis also identified three notable assembly errors resulting in chimeric Z-autosome scaffolds. Cytogenetic analysis further revealed a large W-chromosome that is partially euchromatic, consistent with being a neo-W chromosome. The discovery of a neo-Z and the provisional assignment of chromosome linkage for >90% of *D. plexippus* genes lays the foundation for novel insights concerning sex chromosome evolution in this female-heterogametic model species for functional and evolutionary genomics.

## Background

Major rearrangements of karyotype and chromosome structure often have substantial evolutionary impacts on both the organisms carrying such mutations and the genes linked to such genomic reorganization (Lynch and Walsh 2007; Soltis and Soltis 2012). Additionally, such large-scale chromosomal mutations often present novel opportunities to investigate molecular evolutionary and functional genetic processes, for instance the evolution of neo-sex chromosomes, which can arise from the fusion of an autosome with an existing and well-differentiated allosome. This effectively instantaneous transformation of a formerly autosomal set of genes into sex-linked loci is fertile ground for advancing our understanding of the distinct set of evolutionary forces acting on sex chromosomes relative to autosomes (Bachtrog *et al.* 2009; Pala, Hasselquist, *et al.* 2012; Bachtrog 2013; Šíchová *et al.* 2013). Furthermore, when such an event is observed in a tractable genetic model system, there is opportunity to explore the functional and mechanistic changes associated with sex chromosome evolution. The congruence of neo-sex chromosomes existing in a model system is relatively rare, although there are some notable examples. Namely, independent origins of neo-sex chromosomes are known in *Drosophila* fruit flies (Counterman *et al.* 2004; Flores *et al.* 2008; Bachtrog *et al.* 2009; Zhou *et al.* 2013; Brown and Bachtrog 2014; Nozawa *et al.* 2014) and stickleback fish, where neo-sex chromosomes appear to play an important role in reproductive isolation between incipient species (Kitano *et al.* 2009; Yoshida *et al.* 2014; White *et al.* 2015).

Looking beyond these established model systems, the rapid expansion of genomic technologies has allowed extensive analyses of gene content, sex-biased gene expression, dosage compensation, and sequence divergence for recently evolved sex chromosomes among a very diverse set of organisms including several insect lineages [Teleopsid flies, a grasshopper, and Strepsiptera (Baker and Wilkinson 2010; Mahajan and Bachtrog 2015; Palacios-Gimenez *et al.* 2015)], vertebrates [mammals and birds (Zhou *et al.* 2008; Pala, Hasselquist, *et al.* 2012; Murata *et al.* 2015)], and plants [*Silene* and *Rumex* genera (Hough *et al.* 2014; Charlesworth 2015; Papadopulos *et al.* 2015)]. A clear consensus emerges from this research that the lack of recombination associated with sex chromosomes catalyzes a cascade of evolutionary changes involving the degeneration of one allosome, the accumulation of genes with sex-biased expression, increased evolutionary rates, and often, but not always, the acquisition of dosage compensation. Yet many of the details in this process remain elusive and unresolved, including the rate of allosome divergence, the role of positive selection versus drift, the importance of sex-specific selection, and the mechanisms underlying dosage compensation or the reasons for its absence. It is therefore important to continue identifying new opportunities for novel insight into the evolution of sex chromosomes.

Overwhelmingly, research on sex chromosomes occurs in male-heterogametic (XY) species (Vicoso and Charlesworth 2006; Ellegren 2011; Bachtrog 2013; Parsch and Ellegren 2013). This appears to be particularly true for neo-sex chromosomes, where contemporary genomic analyses of neo-Z or neo-W chromosomes are currently lacking, with one notable exception for birds (Pala, Hasselquist, *et al.* 2012). This imbalance is unfortunate, because ZW sex determination offers the novel framework of female-specific selection during the evolution of heterogamety and is a common form of sex determination in both vertebrates and invertebrates. Birds are the most prominent vertebrate taxon that is female-heterogametic, but it appears that avian neo-sex chromosomes are quite rare and indeed absent from prominent model species, *e.g.* chicken, zebra finch (Nanda *et al.* 2008; Pala, Naurin, *et al.* 2012). Fishes and squamates seem to be far more labile in sex-chromosome constitution, with numerous independent transitions between male and female-heterogamety and relatively frequent sex-autosome fusions (Pennell *et al.* 2015), thus there are potentially great opportunities in these taxa. However, no tractable ZW model system with neo-sex chromosomes has been identified for these lineages.

For many reasons, Lepidoptera, moths and butterflies, may be the most promising female-heterogametic taxon for studying neo-sex chromosomes. Synteny, *i.e.* the chromosomal placement of orthologous genes between species, is unusually well-conserved in Lepidoptera (Pringle *et al.* 2007; The Heliconius Genome Consortium 2012; Ahola *et al.* 2014; Kanost *et al.* 2016), yet there are also numerous known examples of independently evolved neo-Z and neo-W chromosomes, several of which have been well-characterized cytogenetically (Traut *et al.* 2008; Yoshido *et al.* 2011; Nguyen *et al.* 2013; Šíchová *et al.* 2013; Smith *et al.* 2016). Furthermore, comparative genomic resources in this insect order are substantial and growing quickly (www.lepbase.org).

In this context, we report the discovery of a neo-Z chromosome in the monarch butterfly, *Danaus plexippus*, and closely related species. Monarch butterflies, renowned for their annual migration across North America, already have a strong precedent as a model system in ecology (Urquhart 1976; Oberhauser and Solensky 2004). Recently, monarchs have emerged as a model system for genome biology, with a well-assembled reference genome, extensive population resequencing data, and a precedent for genome engineering (Zhan *et al.* 2011, 2014; Merlin *et al.* 2013; Markert *et al.* 2016). The discovery of a neo-Z chromosome further enriches the value of this species as a research model in genome biology and lays the foundation for extensive future insights into the evolution and functional diversity of sex chromosomes.

## Materials and methods

### Sequencing coverage analysis

Illumina shotgun genomic DNA sequencing data for three male and three female *D. plexippus* individuals were selected for analysis from samples sequenced by Zhan *et al.* (2014). Details of sample identities, including GenBank SRA accessions, are given in Supplementary Table S1. Male-female pairs were selected on the basis of approximately equal sequencing coverage. Samples were aligned to the *D. plexippus* version 3 genome assembly with bowtie2 (v2.1.0), using the “very sensitive local” alignment option (Langmead and Salzberg 2012; Zhan and Reppert 2013). The resulting alignments were parsed with the *genomecov* and *groupby* utilities in the BedTools software suite (v2.17.0) to obtain a per-base median coverage depth statistic for each scaffold (Quinlan and Hall 2010). Genomic sequencing data from other *Danaus* species, also generated by Zhan et al. (2014b), were aligned to the same assembly using Stampy (v1.0.22) (default parameters, except for *substitutionrate=0.1*) (Lunter and Goodson 2011).

Coverage analyses comparing males and females were limited to scaffolds of lengths equal to or greater than the N90 scaffold (160,499 bp) (Zhan and Reppert 2013). Also, incomplete cases were excluded (i.e., scaffolds with no reads from one or more samples). In total, 140 scaffolds were excluded, leaving 5,257 scaffolds analyzed. For each sample, each scaffold’s median coverage was divided by the mean across all scaffold median coverages, thereby normalizing for differences in overall sequencing depth between samples. Samples were grouped by sex and the per-scaffold mean of normalized coverage depth was compared between sexes, formulated as the log_2_ of the male:female coverage ratio. Autosomal scaffolds are expected to exhibit equal coverage between sexes, yielding a log_2_ ratio of zero. Z-linked scaffolds should have a ratio of one, due to the two-fold greater representation in males. Manipulation, analysis, and visualization of coverage data was performed in R (R Developement Core Team 2015).

To select scaffolds with intermediate median coverage ratios, we used Bedtools *genomecov* to calculate per-base coverage, in order to identify potential assembly errors producing Z-Autosomal chimeric scaffolds. For each sample, coverage per base was divided by the mean of all scaffold median coverage values, thus normalizing for overall sequencing depth. The normalized coverage per base was averaged within sex and visualized along the length of the scaffold by using the median of a 5kbp sliding window, shifted by 1kbp steps.

Point estimates for Z-autosomal break points in chimeric scaffolds were generated using a sliding window analysis of male:female coverage ratios. Putative break points were obtained as the maximum of the absolute difference between adjacent non-overlapping windows. A window of 150 Kbp with 10 kbp steps was used for DPSCF300001 and the 5’ break point of DPSCF30028. A window of 10 kbp with 1 kbp steps was used for DPSCF30044 and in a second, localized analysis between 1.5 Mb and the 3’ terminus of DPSCF30028 to localize the second, 3’ break point.

### Orthology-based chromosomal assignments for *D. plexippus* scaffolds

Putative chromosomal linkage was predicted for *D. plexippus* scaffolds relative to the genome assemblies of three reference species, *M. cinxia, B. mori,* and *H. melpomene* (the Glanville fritillary, domestic silkmoth, and postman butterfly), based on counts of orthologous genes (The International Silkworm Genome Consortium 2008; The Heliconius Genome Consortium 2012; Ahola *et al.* 2014). Orthologous proteins were predicted between *D. plexippus* and each reference species using the *Proteinortho* pipeline (Lechner *et al.* 2011). Using only 1-to-1 orthologs, we tabulated per *D. plexippus* scaffold the number of genes mapped to each chromosome in the reference species. Each *D. plexippus* scaffold was assigned to the chromosome with the highest count of orthologs in the reference species. Scaffolds were excluded from analysis when maximum ortholog count was tied between two or more scaffolds, though this situation was rare and involved scaffolds with low genes counts.

### Point estimate of the Z-autosome fusion

The fusion point in Monarch between ancestrally Z and autosomal segments was localized by aligning the homologous *H. melpomene* or *M. cinxia* chromosomes against Monarch scaffold DPSCF300001 (Ahola *et al.* 2014; Davey *et al.* 2016). Alignments were based on six-frame amino acid translations using the PROmer algorithm and visualized with mummerplot, both from the MUMmer software package (v3.1) (Kurtz *et al.* 2004). We initially aligned the complete set of scaffolds from the Z (HmChr21, McChr1) or relevant autosome (HmChr2, McChr21), yielding a preliminary indication that the Z-autosome fusion point occurred at ∼4 Mbp on DPSCF300001. To refine and better visualize this phenomenon, pseudo-assemblies were created for each chromosome using query scaffolds producing >500 bp of total aligned coverage on DPSCF300001. Selected query scaffolds were concatenated into a single fasta entry, with ordering based on target alignment positions. For each species, the Z and autosomal pseudo-assemblies were co-aligned to DPSCF300001. The transition point between contiguous alignments of the two pseudo-assemblies from distinct chromosomes was interpreted as the approximate location of the Z-autosome fusion in Monarch.

### Cytogenetic analysis

All *D. plexippus* tissues used for cytogenetic analysis were from captive-bred butterflies reared on an artificial diet provided by MonarchWatch (MonarchWatch.org). Spread chromosome preparations were made from gonads of third to fifth instar larvae of both sexes following Mediouni *et al.* (2004). In order to test for the presence of sex chromatin, preparations of polyploid somatic nuclei were made according to Traut *et al.* (1986) from Malpighian tubules dissected from the same material.

Genomic DNA was isolated separately from males and females by standard phenol-chloroform extraction. Briefly, larval tissues were homogenized in liquid nitrogen, transferred in lysis buffer (100 mM NaCl, 10 mM Tris-HCl pH 8,0, 50 mM EDTA, 100 μg/ml Proteinase K, 0,5% Sarkosyl), and incubated at 37°C overnight. The samples were then treated with RNase A (10 µg/ml) and purified by three phenol, one phenol-chloroform, and one chloroform extractions. Male- and female-derived hybridization probes were labeled by nick translation as described in Šíchová et al. (2015).

Genomic in situ hybridization (GISH) was performed as described by Fuková et al. (2005). Comparative genomic hybridization (CGH) was carried out according to the protocol in Šíchová et al. (2013) with several modifications, as follows. Prior to denaturation, RNase A treated slides were incubated in 5x Denhardt’s solution (0,1% Ficoll, 0,1% polyvinylpyrrolidone, 0,1% bovine serum albumin) at 37°C for 30 min. A 10 µl hybridization mixture consisting of labeled female and male probes (350 ng each), sonicated salmon sperm DNA (25 µg), 50% deionized formamide, and 10% dextran sulfate in 2x SSC was denatured and allowed to reanneal at 37°C for two hours (c.f. Kallioniemi *et al.* 1992) before it was hybridized to the denatured female preparation.

Results were documented in a Zeiss Axioplan 2 microscope (Carl Zeiss, Jena, Germany) equipped with appropriate fluorescence filter set. Images were captured with an Olympus CCD monochrome camera XM10 equipped with cellSens 1.9 digital imaging software. The images were pseudocoloured and superimposed with Adobe Photoshop CS3.

### Data Availability

Putative chromosomal assignments for D. plexippus genes are provided in Supplemental Table S2. Estimated breakpoints reported for chimeric Z-autosomal assemblies are provided in Supplemental Table S3. Other intermediate Rresults files and code used in described analyses are available upon request.

## Results

### Identifying Z-linked scaffolds in *D. plexippus*

We identified Z-linked scaffolds in the *D. plexippus* genome assembly (Zhan *et al.* 2011; Zhan and Reppert 2013) by comparing sequencing coverage from male and female samples. Males should have twice the Z chromosome DNA content than females, while autosomes should have equal DNA content between sexes. Thus a corresponding two-fold difference in sequencing coverage is expected between sexes for the Z chromosome, but not autosomes, and can be used to identify Z-linked scaffolds (Martin *et al.* 2013; Vicoso *et al.* 2013; Mahajan and Bachtrog 2015). A histogram of male:female ratios of median coverage clearly identifies two groups of scaffolds (Fig. 1). One large cluster is centered around equal coverage between sexes (Log_2_ M:F = 0) and a second, smaller cluster is centered around two-fold greater coverage in males (Log2 M:F =1). We can thus clearly distinguish the Z-linked scaffolds as those with Log_2_(M:F) > 0.5, with the remainder of the scaffolds presumed to be autosomal.

**Figure 1.**
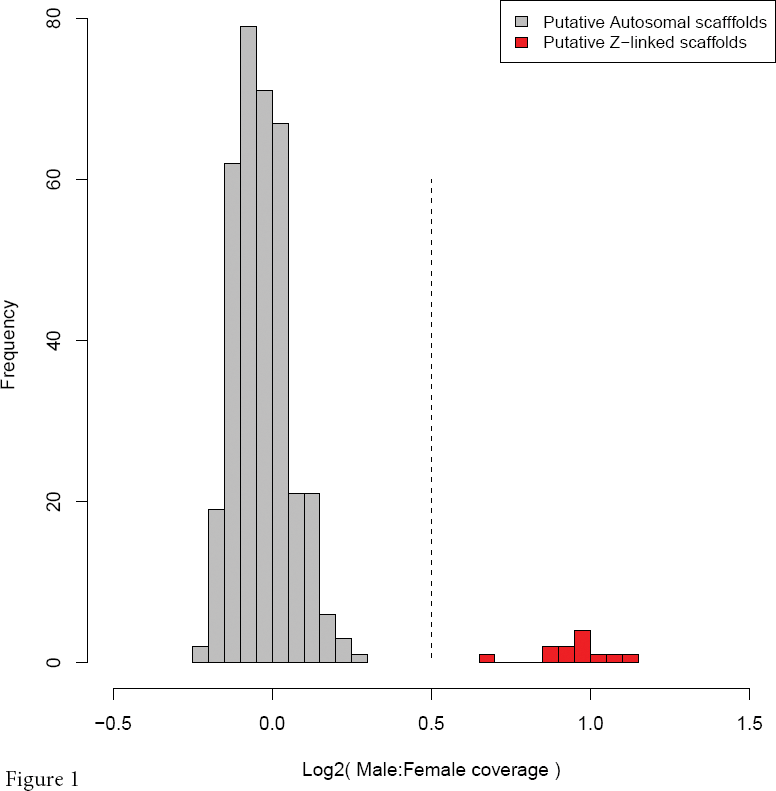
Distribution of median normalized male:female genomic sequencing coverage ratios for *D. plexippus* version 3 assembly scaffolds. Only scaffolds of length equal to or greater than the N90 scaffold are shown. The dotted line at 0.5 represents the value used to partition scaffolds as autosomal (grey) or Z-linked (red).

One scaffold, DPSCF300028, appeared to have an intermediate coverage ratio, falling at Log_2_ M:F ≈ 0.7. One likely explanation for such an intermediate value is that the scaffold is a chimera of Z-linked and autosomal sequence arising from an error in genome assembly (Martin *et al.* 2013). In this scenario, only a portion of the scaffold is Z-linked and gives a two-fold difference in coverage between sexes; the remaining autosomal fraction of the scaffold yields equal coverage. The resulting estimate of average coverage for the entire scaffold then falls at a value between expectations for Z or autosomal scaffolds. This is clearly true for DPSCF300028, as revealed by examining basepair-level sequencing coverage across the scaffold (Fig. 2A). While average male coverage is consistent across the entire length of the scaffold, female coverage exhibits a clear transition between coverage equal to males (the autosomal portion) and coverage one half that of males (the Z-linked portion). Indeed, there are two such transitions in scaffold DPSCF300028, which we estimate to occur at 0.76 Mbp and 1.805 Mbp, creating a Z segment flanked by autosomal segments.

**Figure 2.**
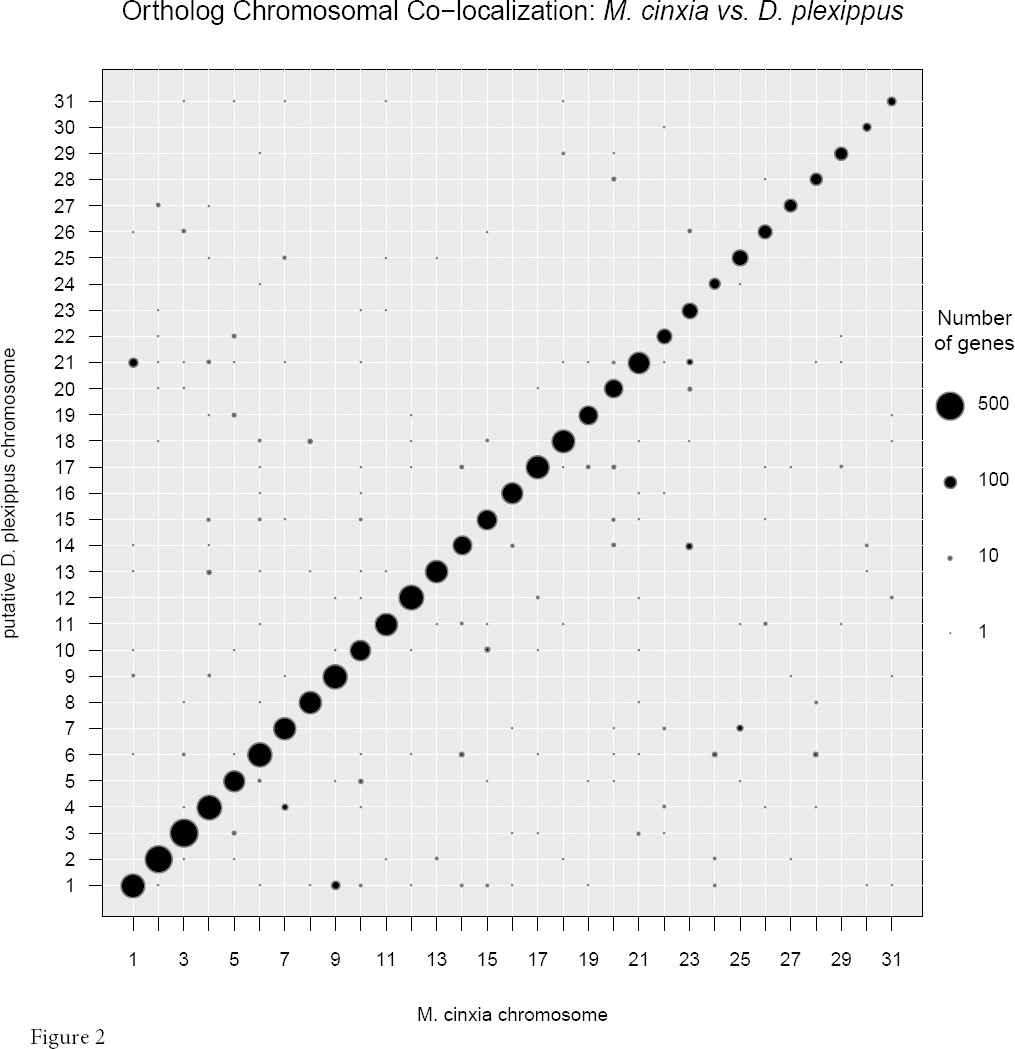
Normalized male and female coverage along the length of chimeric scaffolds, for (A) DPSCF300028, (B) DPSCF300044, and (C) DPSCF300001. Coverages are plotted as sliding windows (width = 5Kbp, step = 1 Kbp) of median basepair values. The associated male:female ratio of coverage for each window is plotted as a red line below the pair of sex-specific plots. Asterisks indicate the estimated break point between Z linked and autosomal segments of each scaffold, as determined by the maximum difference in adjacent, non-overlapping windows of male:female ratio (see methods for details).

### Ortholog counts link scaffolds to chromosomes

As mentioned above, Lepidoptera show a very high level of synteny conserved across substantial evolutionary distances (Pringle *et al.* 2007; The Heliconius Genome Consortium 2012; Ahola *et al.* 2014; Kanost *et al.* 2016). Thus it is possible to use counts of orthologous genes to assign *D. plexippus* scaffolds to linkage groups (*i.e.* chromosomes) delineated in other moth or butterfly species. We generated predicted orthologs between *D. plexippus* and three other reference species where genetic linkage mapping has been used to assign genomic scaffolds to chromosomes: *Melitaea cinxia* (N=31), *Heliconius melpomene* (N=21), and *Bombyx mori* (N=28) (The International Silkworm Genome Consortium 2008; The Heliconius Genome Consortium 2012; Ahola *et al.* 2014). *M. cinxia* and *H. melpomene* are both butterflies equally diverged from *D. plexippus* and all three are part of the same family, Nymphalidae; the silkmoth, *B. mori*, is distinctly more diverged, located outside of the suborder containing all butterflies (Wahlberg *et al.* 2009; Kawahara and Breinholt 2014).

To assign *D. plexippus* scaffolds to chromosomes, we tabulated per scaffold the counts of one-to-one reference species orthologs per reference species chromosome. *D. plexippus* scaffolds were then assigned to the reference chromosome with the maximum count of orthologs. For a few scaffolds, a tie occurred in maximum ortholog count per reference chromosome, in which case the scaffold was removed from further analysis; at most this occurred for only 14 scaffolds per reference species and usually involved small scaffolds harboring fewer than 5 orthologs. Typically, this method yielded a single obvious reference chromosomal assignment for each *D. plexippus* scaffold.

This method of ortholog-count chromosomal lift-over resulted in putative chromosomal assignments for >90% of *D. plexippus* genes relative to each reference species (Table 1, Supplementary Table S2). Also, at least 4500 orthologous genes were co-localized to chromosome between *D. plexippus* and each reference species. Having several thousand orthologs mapped to chromosome in *D. plexippus* and a reference species presents the opportunity to examine the extent of chromosomal rearrangements and gene movement between the two species. Here we primarily report the comparison with *M. cinxia* because this species is believed to retain the ancestral lepidopteran karyotype of 31 chromosomes (Ahola *et al.* 2014). Furthermore, this count of chromosomes is closest to that reported for several *Danaus* butterflies, including monarch (N=30, see Figure 3), making it the most appropriate comparison available (Brown *et al.* 2004). *H. melpomene* and *B. mori* are known to have less similar karyotypes involving several chromosomal fusions relative to *M. cinxia*; nonetheless, details of comparisons to these two species are reported in the supplementary content and provide comparable support for the primary findings reported here.

**Table 1.** Table 1. Summary of assigning *D. plexippus* genes and scaffolds to chromosomes via orthology “liftover” relative to *M. cinxia*.

|  |  |
| --- | --- |
| 1:1 orthologs identified | 6,740 |
| 1:1 orthologs assigned to *M. cinxia* chromosome | 4,607 (68.4%) |
| Protein coding genes in *D. plexippus* | 15,130 |
| *D. plexippus* protein coding genes assigned to chromosome | 14,129 (93.4%) |
| *D. plexippus* scaffolds putatively assigned to chromosomes | 454 |

**Figure 3.**
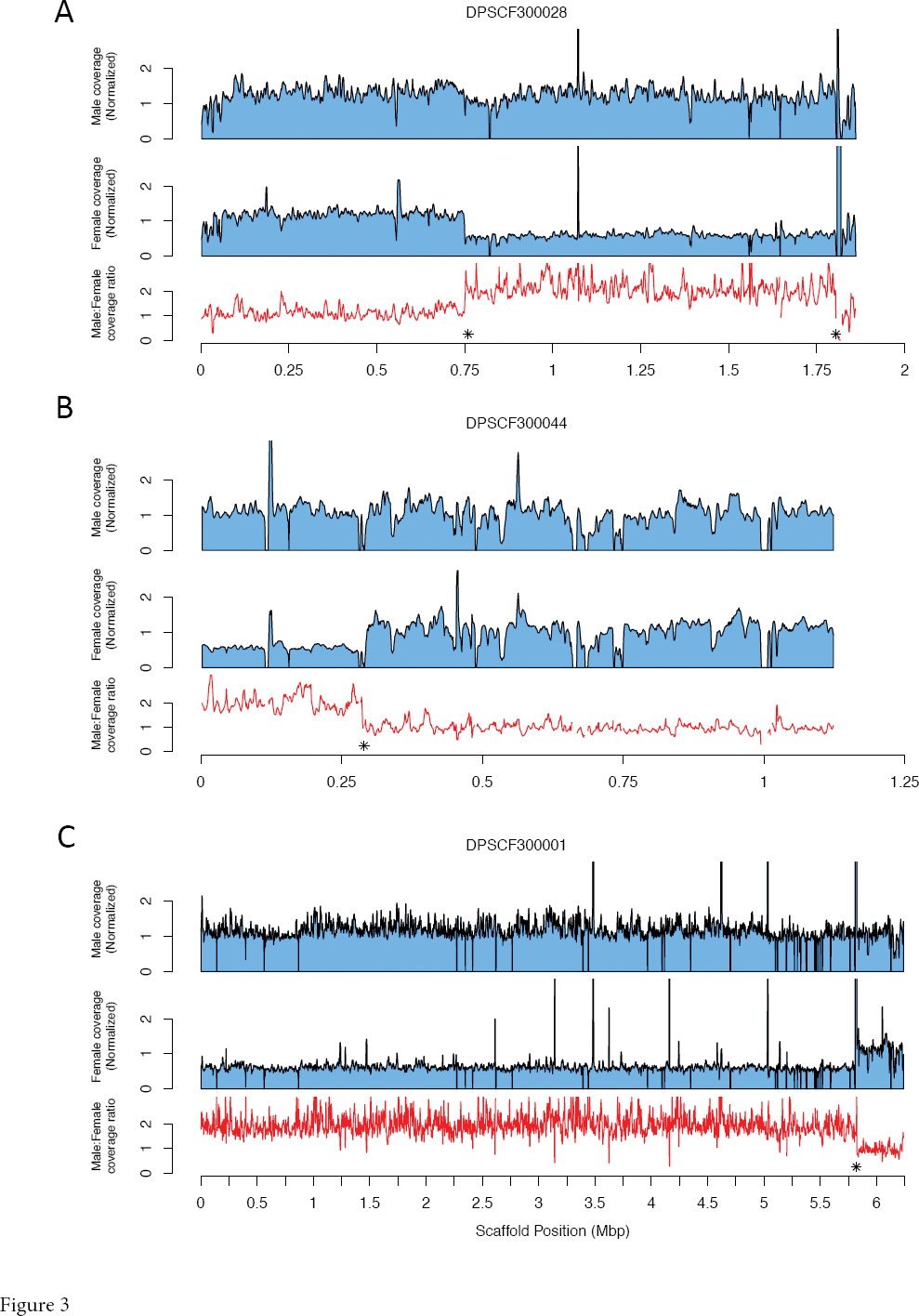
Chromosomal co-linkage between *D. plexippus* and *M. cinxia* for predicted orthologous proteins.

Figure 3 summarizes the cross-tabulation of chromosomal linkage for >4500 orthologs between *M. cinxia* and *D. plexippus*. The overwhelming majority of orthologs fall on the diagonal, indicating substantial conservation of chromosomal linkage and relatively little gene shuffling, as has been reported elsewhere for Lepidoptera (The Heliconius Genome Consortium 2012; Ahola *et al.* 2014; Kanost *et al.* 2016). The two most notable exceptions to this pattern both involve the Z chromosome (Chr1). In one case [McChr9, DpChr1] we could anticipate this because of the previously identified chimeric scaffold, DPSCF300028. This scaffold harbors 34 orthologs assigned to McChr1 and 23 orthologs assigned to McChr9, consistent with the chimeric nature of the scaffold revealed from male:female coverage ratios (Fig 2A).

The second case [McChr1, DpChr21] appeared to arise entirely from a single scaffold, DPSCF300001, the largest scaffold in the *D. plexippus* v3 assembly. This scaffold carried 107 orthologs assigned to McChr21, 28 orthologs assigned to McChr1, 13 orthologs assigned to McChr23, and a few other orthologs assigned to other autosomes. Notably, despite the large number of apparently autosomal orthologs, the average male:female coverage ratio for DPSCF300001 was consistent with it being Z-linked [ Log_2_(M:F coverage) = 0.92]. Nonetheless, we plotted coverage across the chromosome and detected a ∼1 Mbp portion at the 3’ end of the scaffold with coverage patterns consistent with being an autosome (Fig 2C). The *M. cinxia* orthologs in this autosomal portion, with an estimated breakpoint at 5.82 Mbp, were linked exclusively to McChr23. There was not an obvious shift in sequencing coverage between sexes to indicate a misassembled Z-autosome chimera involving McChr21. Rather, it appeared that nearly the entirety of scaffold DPSCF300001 had twice the coverage in males than in females, consistent with Z-linkage for regions apparently homologous both to Mc1(Z) and McChr21.

### A neo-Z chromosome in *D. plexippus*

The observation that a substantial portion of scaffold DPSCF300001 was Z-linked and homologous to McChr21, while another large section of the same scaffold was homologous to McChr1, i.e. McChrZ, led us to hypothesize that a Z-autosome fusion could readily explain the karyotypic differences between *D. plexippus* (N=30) and *M. cinxia* (N=31). To further investigate this hypothesis of a major evolutionary transition in sex chromosome composition in the *Danaus* lineage, we examined the chromosomal assignments for all Monarch scaffolds identified as Z-linked via sequencing coverage ratios (Z-cov scaffolds). Specifically, we identified the unique set of reference chromosomes to which Z-cov scaffolds were assigned, and then examined the male:female coverage ratio for all scaffolds assigned to those reference chromosomes. In the case of *M. cinxia* as the reference, all Z-cov scaffolds were assigned either to McChr1 or McChr21 (Fig. 4; comparable results were obtained for *H. melpomene* and *B. mori*, Supplementary Fig. S2). This result provides further evidence that the Z in *D. plexippus* is a neo-sex chromosome reflecting the fusion of the ancestral Z chromosome with an autosome homologous to McChr21.

**Figure 4.**
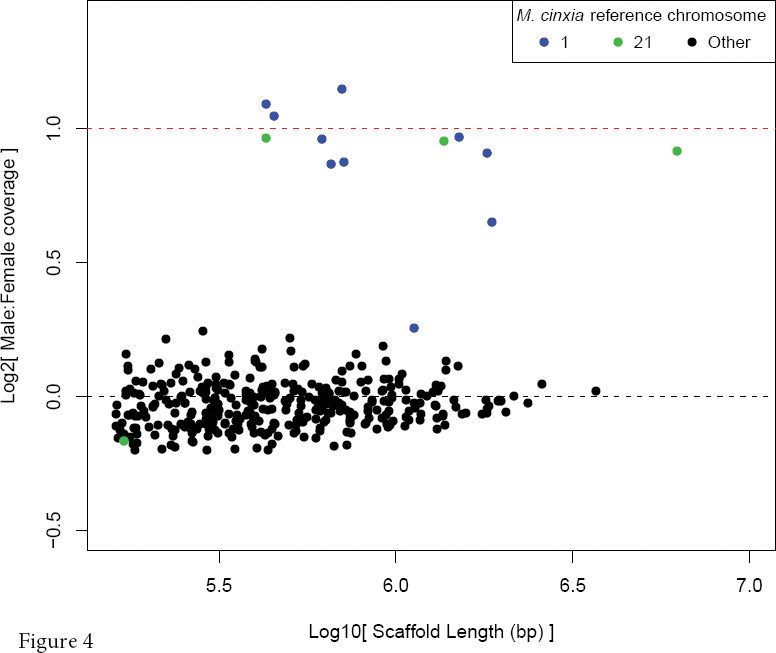
Ratios of male:female median normalized genomic sequencing coverage plotted against scaffold length. Scaffolds homologous via “liftover” procedure to *M. cinxia* chromosomes 1/Z (blue) and 21 (green) are plotted in distinct colors. Dotted lines indicate expected values for Z-linked (red) and autosomal (black) scaffolds.

This analysis intersecting Z-cov scaffolds with homology to *M. cinxia* revealed two scaffolds that did not fit with the expected pattern of sequencing coverage (Fig. 5). First, scaffold DPSCAF300044 was assigned to McChr1(Z) but had Log_2_ M:F ≈ 0.25, much more like other autosomes than other Z-linked chromosomes. This scaffold had seven Z-linked orthologs and four autosomal, suggesting another chimeric scaffold. Indeed, examining coverage across the scaffold revealed a clear transition in coverage as previously observed for DPSCF300001 and DPSCF300028 (Fig 2B). Thus the low male:female coverage ratio for this scaffold is likely the artifact of an assembly error. Again we were able to partition the scaffold into two sections, one autosomal and one Z-linked, with a breakpoint estimated at 0.29 Mbp from the 5’ end. The autosomal section contained approximately equal counts of orthologs assigned to two distinct chromosomes in *M. cinxia* and the other reference species, so linkage to a specific autosome could not be predicted. Supplementary Table S3 summarizes breakpoints and predicted scaffold assignments for the three chimeric Z-autosome scaffolds identified here.

**Figure 5.**
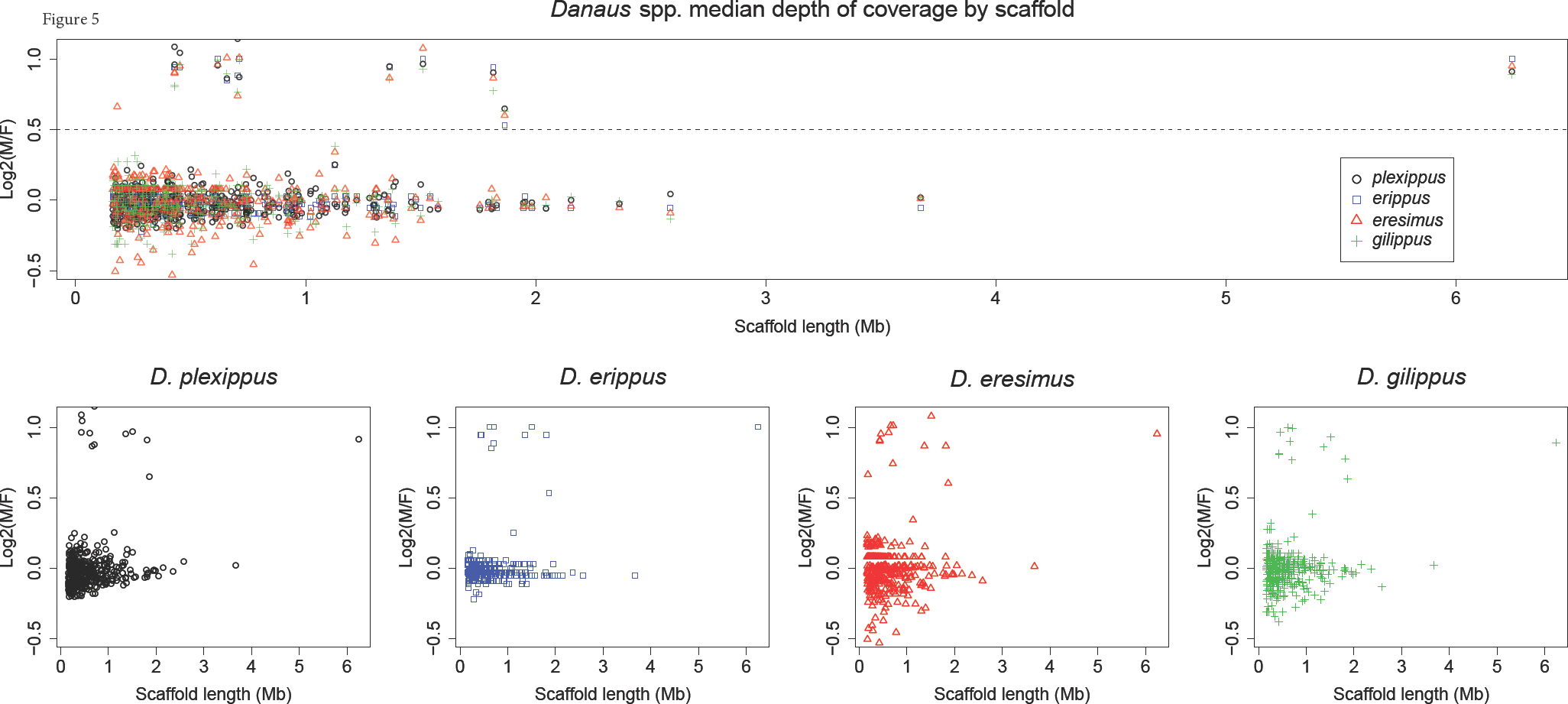
Ratios of male:female median normalized genomic sequencing coverage plotted by scaffold length for four species of *Danaus* butterflies. The dotted line at log_2_(M/F)=0.5 represents the threshold used to discern autosomal (<0.5) from Z-linked (>0.5) scaffolds.

DPSCF300403 was the other scaffold where the M:F ratio of median coverage was inconsistent with the hypothesis of a neo-Z chromosome. This scaffold was assigned to McChr21 but had an autosomal coverage ratio. Coverage along the chromosome was consistent with it being entirely autosomal (Supplementary Figure S3). In this case the scaffold carried only a single one-to-one orthologous gene (and only 5 protein-coding genes total), so the assignment to McChr21 is tenuous and likely inaccurate. This scaffold also had a single one-to-one ortholog found in *B. mori*, and none identified in *H. melpomene*. We therefore consider this scaffold largely uninformative concerning the presence of a neo-Z in *D. plexippus*.

### The neo-Z chromosome exists in the Monarch’s close relatives

The Monarch population genomic dataset from Zhan et al. (2014b) also contained male and female resequencing samples from three closely related congeners: *D. gilippus, D. erippus,* and *D. eresimus*. This allowed us to assess whether this neo-Z exists in these related species similarly to *D. plexippus*. Analyzing male versus female resequencing in these species does indeed show the same scaffolds homologous to both McChr1 and McChr21 as having coverage differences consistent with a neo-Z (Fig. 5). Thus it appears that the origin of this neo-Z predates the diversification of the genus *Danaus*.

### Annotating chromosomal linkage

The combination of sequencing coverage analysis and comparative lift-over allowed us to provisionally assign most genes to chromosomes in *D. plexippus*. Genes falling on Z-cov scaffolds, or within the portion assessed as Z-linked for noted chimeric scaffolds, have been assigned to the Z chromosome. We further partitioned these Z-linked genes into being on the ancestral (anc-Z) or neo (neo-Z) portion of the Z, based on scaffold homology to reference chromosomes. In the case of DPSCF300001, we localized the fusion point between anc-Z and neo-Z by aligning *M. cinxia* and *H. melpomene* scaffolds from the Z (HmChr21, McChr1) or relevant autosome (HmChr2, McChr21). Alignments with both species were consistent in placing the fusion point at approximately 3.88 Mbp from the 5’ end of the scaffold (Supplementary Fig. S5). Otherwise, genes and scaffolds were assigned to chromosomes based directly on the results of the lift-over relative to *M. cinxia*. Table 2 gives a tabulated summary of results, while results for every protein coding gene are provided in Supplementary Table S4.

**Table 2.** Summary of provisional chromosomal linkage for *D. plexippus* protein coding genes, with chromosomal identity reflecting homology to *M. cinxia*

| Chromosome | Number of Genes |
| --- | --- |
| 1 | 1101 |
| Anc-Z | 624 |
| Neo-Z | 477 |
| 2 | 704 |
| 3 | 758 |
| 4 | 582 |
| 5 | 494 |
| 6 | 689 |
| 7 | 483 |
| 8 | 535 |
| 9 | 647 |
| 10 | 452 |
| 11 | 574 |
| 12 | 576 |
| 13 | 501 |
| 14 | 429 |
| 15 | 493 |
| 16 | 524 |
| 17 | 604 |
| 18 | 561 |
| 19 | 414 |
| 20 | 399 |
| 22 | 302 |
| 23 | 318 |
| 24 | 185 |
| 25 | 329 |
| 26 | 274 |
| 27 | 250 |
| 28 | 284 |
| 29 | 294 |
| 30 | 151 |
| 31 | 168 |
| Not Assigned | 1055 |

### Cytogenetic analysis

Preparations of highly polyploid nuclei of Malpighian tubules from *D. plexippus* larvae were examined for the presence of a so-called “sex chromatin”, *i.e.* female specific heterochromatin body consisting of multiple copies of the W chromosome (Traut and Marec 1996). Large multi-lobed nuclei were observed on both male and female preparations (Fig. 6), which suggests a high degree of polyploidy in the examined cells, as expected in Malpighian tubules (cf. Buntrock *et al.* 2012). All female nuclei contained a single, highly stained heterochromatin body (Fig. 6a). In contrast, no such heterochromatin was detected in male somatic nuclei (Fig. 6b). This discrepancy confirms the female-specificity of the heterochromatin observed and indicates a W chromosome is a component of the *D. plexippus* genome.

**Figure 6.**
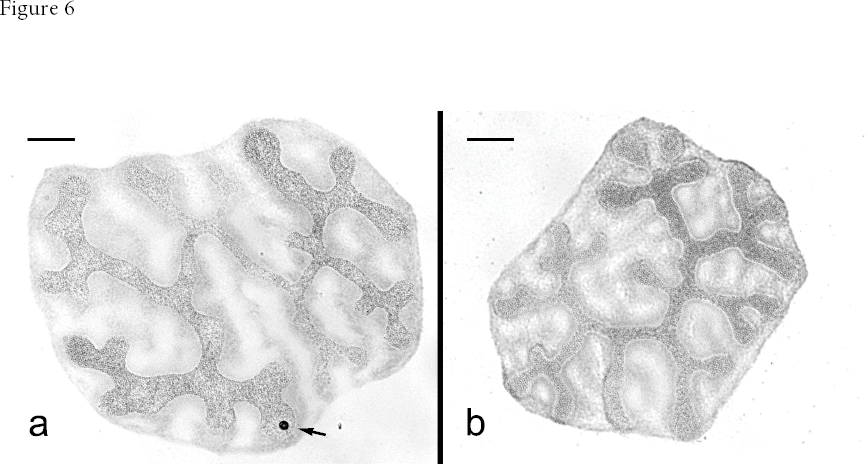
Multi-lobed, highly polyploid nuclei of the Malpighian tubules from *Danaus plexippus* larvae stained by orcein. (a) A female nucleus with a single deeply stained sex chromatin body (arrow). (b) A male polyploid nucleus with no heterochromatin. Bars = 20 µm.

Spread mitotic complements of males contained 2N=60 chromosomes. The chromosomes were generally small and uniform, as is typical of lepidopteran karyotypes (Marec *et al.* 2009), except for two distinctly larger elements (Fig. 7a). Female mitotic metaphase consisted of 2N=60 elements as well. In females, however, the two largest chromosomes differ in intensity of their DAPI staining. The deeply stained element found exclusively in females presumably represents the W sex chromosome consisting of A-T rich heterochromatin (Fig. 2b; Kapuscinski 1979).

**Figure 7.**
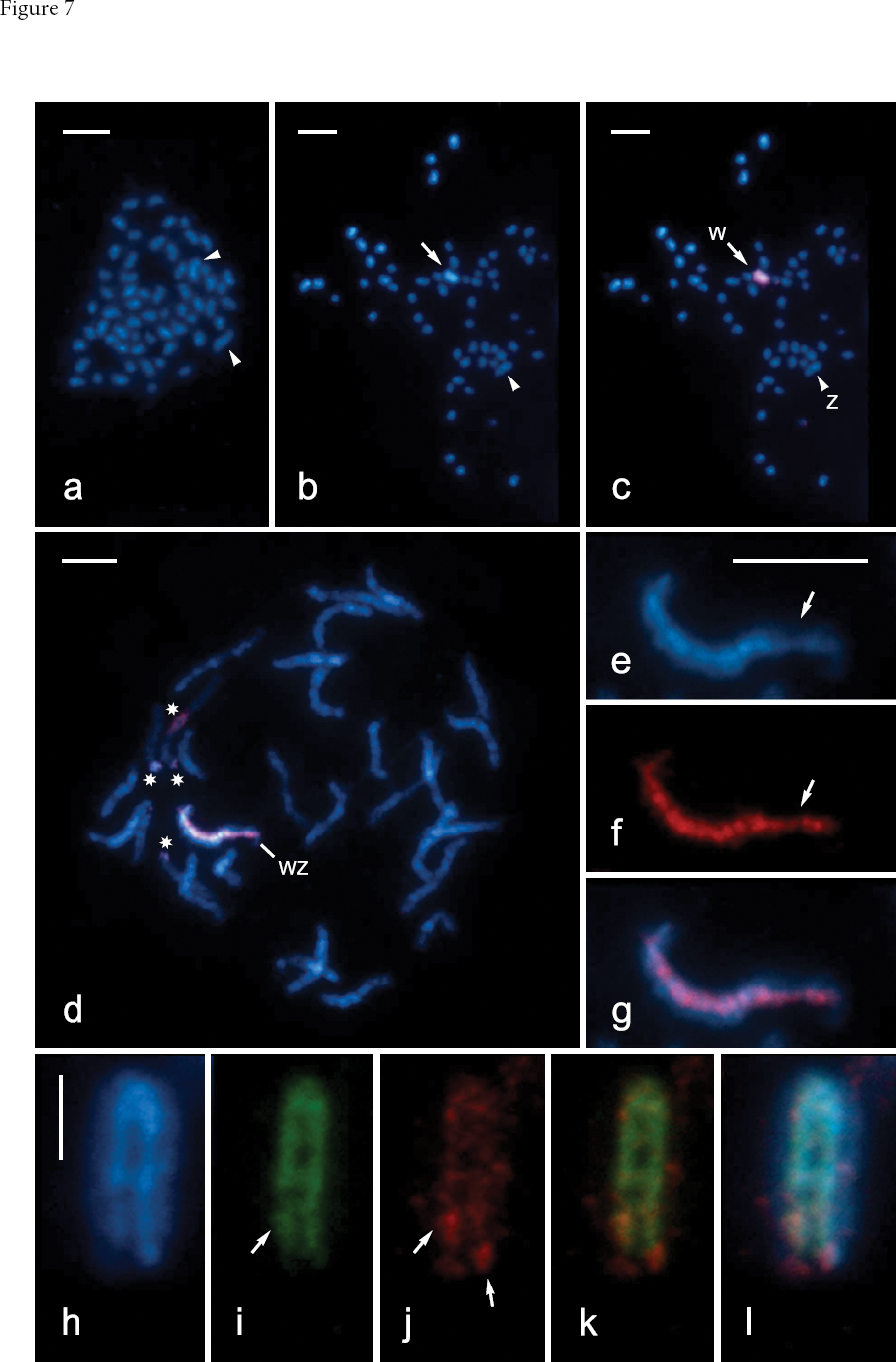
Cytogenetic analysis of sex chromosomes in *D. plexippus*. Chromosomes were counterstained by DAPI (blue). **(a)** Male mitotic metaphase complement consisting of 2N=60 elements. Note the large chromosome pair (arrowheads). **(b)** Female mitotic metaphase nucleus comprising 2N=60 chromosomes. The largest elements differ in intensity of their staining (arrowhead and arrow, the latter marking highly stained chromosome). **(c-g)** Female complements examined by genomic in situ hybridization (GISH). Female derived probe was labeled by Cy3 (red). **(c)** The same mitotic complement as in (b). The probe identified the DAPI positive chromosome as a W chromosome and thus indicated the other large chromosome to be the Z chromosome. **(d)** Female pachytene complement consisting of 30 bivalents. The WZ bivalent was clearly recognized by female-derived probe, which highlighted whole W chromosome except its terminal segments. The probe also marked three terminal and one interstitial regions of several autosomes (asterisks). **(e)** The WZ bivalent stained by DAPI. Note that only about two thirds of the W chromosome thread is deeply stained and apparently heterochromatic. The more lightly stained euchromatic segment is marked by an arrow. **(f)** Hybridization signal of female-derived probe highlighting the W chromosome thread. Note that the signal is weaker in the euchromatic region (arrow). **(g)** Composite image of DAPI and the probe. **(h-l)** A pachytene WZ bivalent probed by comparative genomic hybridization (CGH). Male-derived probe was labeled by Cy3 (red), female-derived probe by fluorescein (green). **(h)** The WZ bivalent stained by DAPI. The weakly stained Z is wrapped around the strongly stained W, which is folded in half. **(i)** Hybridization signal of female-derived probe. Note that the probe labeled only a small region of the euchromatic segment (arrow). **(j)** Hybridization signal of male-derived probe strongly highlights two W segments (arrows). **(k)** Both the female- and male-derived signals merged. **(l)** Composite image of DAPI and both probes. Bars = 5 µm in (a-g) and 2.5 µm in (h); (e-g) and (h-l) have the same scale.

In both mitotic and pachytene nuclei, GISH clearly identified the W chromosome by strong binding of female-derived probe (Fig. 7 c, d). It confirmed that the W chromosome is one of the two exceptionally large chromosomes in the *D. plexippus* karyotype (Fig. 7 c). In pachytene oocytes, the WZ bivalent was easily discernible by the heterochromatic W chromosome thread. However, about one third of the chromosome was not strongly highlighted by DAPI and the intensity of its staining was comparable to autosomes, indicating a portion of the W that is substantially euchromatic (Fig. 7 e). Accordingly, the female-derived probe did not highlight the W chromosome homogeneously as the signal was weaker on both its ends and the euchromatic segment (Fig. 7 e-f). Female-derived probes in GISH also strongly labeled one interstitial and a few terminal regions of some autosomes, which most likely contain clusters of repetitive sequences (Fig. 7 d).

Comparative genomic hybridization with both male- and female-derived probes was also used to assess the broad molecular composition of the *D. plexippus* W chromosome (Fig. 7 h-i). Hybridization signal of female-derived probe labeled by fluorescein was largely consistent with the results obtained by GISH. The signal highlighted nearly the entire W chromosome thread, with the exception of its termini and euchromatic segment, in which the probe detected only a small interstitial block (Fig. 7 i, k, l). The male-derived probe labeled by Cy3 provided relatively weak hybridization signal, which was scattered along the W chromosome. This male probe highlighted only two regions: the W chromosome end opposite to the euchromatic segment and the region highlighted within the euchromatin by female-derived probe (Fig. 7 j, k, l). Both probes detected the same autosomal regions as GISH (data not shown).

## Discussion

Using a combination of genomic resequencing, comparative genomics, and cytogenetic analysis, we have documented the presence of a neo-Z chromosome in *Danaus* butterflies, along with what is likely an accompanying neo-W chromosome. This discovery of neo-sex chromosomes in *Danaus* butterflies and our discrimination of genes falling on the ancestral versus recently autosomal portions of the Z are fundamental observations that provide the foundation for a host of future inferences. These results create novel opportunities to address rates of molecular evolution, the evolution of dosage compensation, the pattern of allosome divergence, and many other important questions in sex chromosome biology, all in a female-heterogametic species that is also an emerging genomic model system.

In analyzing patterns of chromosomal fusion in *H. melpomene* and *B. mori* relative to *M. cinxia*, Ahola et al. (2014) report a significant tendency for a limited set of ancestral chromosomes – particularly the smallest ones – to be involved in chromosomal fusion events. Neither the ancestral Z nor McChr21 are among these small, repeatedly fused chromosomes; thus the chromosomal fusion reported here does not fit neatly with this pattern. Nonetheless, HmChr2 (homologous to McChr21) is the second smallest chromosome that remains unfused between these lineages (Davey *et al.* 2016). So it is also difficult to argue strongly that this Z-autosome fusion in *Danaus* is a striking contrast to the trend of chromosomal fusions involving small chromosomes.

Motivated by the bioinformatic discovery of a neo-Z chromosome, we performed cytogenetic analysis of the *D. plexippus* karyotype in order to provide further insight into evolution and molecular composition of the monarch sex chromosomes. Previously, an observation of N=30 chromosomes was reported only for males (Nageswara-Rao and Murty 1975). Our current analysis confirms the same chromosome number not only in males but also in females (Fig. 7a, b). Equal numbers of chromosomes in males and females, along with presence of sex chromatin in females, indicates that a single W chromosome persists in this species alongside the neo-Z. Furthermore, detailed analysis of mitotic complements revealed a large chromosome pair (Fig. 7a, b) and GISH clearly identified one chromosome of the pair as the W chromosome (Fig. 7c). A similar, extraordinarily large chromosome pair was recently shown to correspond to neo-sex chromosomes in leafroller moths of the family Tortricidae (Nguyen *et al.* 2013; Šíchová *et al.* 2013).

GISH represents a simplified version of CGH, which has been successfully used for evaluating the gross molecular composition of lepidopteran W chromosomes (e.g. Mediouni *et al.* 2004; Fuková *et al.* 2005). Previous studies in several moth species contrasted fluorescence intensities of male versus female derived probes and identified two common types of repeats on lepidopteran W chromosomes: (i) repetitive sequences common to both males and females, *i.e.* present in autosomes and Z chromosome; and (ii) repetitive sequences exclusively or predominantly present in females (Sahara *et al.* 2003). In these previously studied species, the W primarily contains the first type, *i.e.* ubiquitous repeats (*e.g.* Fuková *et al.* 2005; Šíchová *et al.* 2013). In contrast, the monarch W appears distinct from the W chromosomes of these other species because the majority of the monarch W chromosome overwhelmingly contains repeats of the second type, *i.e.* female-limited repeats. The monarch W was labelled primarily by female-derived probe (7 i-k), indicating that it is primarily comprised of repetitive sequences either specific to or greatly enriched on the W chromosome. Only two small segments of the W showed notably high densities of ubiquitous repeats commonly enriched on the entire W in other lepidopteran species.

This discrepancy between Monarch and other species could be related to the relatively small size of the *D. plexippus* genome. The monarch butterfly represents the smallest lepidopteran genome yet sequenced, with haploid nuclear content of male 284 Mbp (see Dolezel *et al.* 2003 for the conversion of pg of DNA to Mbp; Gregory and Hebert 2003) and female 273 Mbp (Zhan *et al.* 2011). This small genome is presumably depleted of repetitive sequences found more ubiquitously among the autosomes and Z in other lepidoptera. Indeed, repeat content constitutes only 13.1% of the *D. plexippus* genome assembly (Zhan *et al.* 2011). The results of CGH thus could reflect the fact that the W chromosome represents the last refuge for many such repetitive sequences in the monarch genome after otherwise being purged from the genome.

Cytogenetic analyses thus confirm that the *D. plexippus* W chromosome is well differentiated relative to the Z, which is in agreement with sequencing data that yield a very consistent 2:1 coverage ratio on scaffold regions corresponding to McChr21. If a neo-W retained substantially close homology to the neo-Z, we would expect many sequencing reads emanating from the neo-W to align to the neo-Z, and shift this coverage ratio towards one. This evidently does not occur, indicating substantial divergence between the neo-Z and any neo-W sequence that is retained.

Cytogenetic analyses further indicate that the monarch W chromosome exhibits notable compositional heterogeneity. Both GISH and CGH revealed terminal gaps in female-derived signal (Fig. 2g, l). This is similar to GISH results obtained in a codling moth, *Cydia pomonella* (Tortricidae), where female derived probe labelled the entire W chromosome except for both subtelomeric segments (Fuková *et al.* 2005). Van’t Hof *et al.* (2012) proposed that a copy of the Z-linked *laminin A* gene was transferred and maintained to a W chromosome of the peppered moth, *Biston betularia* (Geometridae), by gene conversion resulting from ectopic recombination between repeats localized in terminal chromosome regions. The same mechanism could be invoked to explain the lack of female specific signals on the monarch W chromosome ends.

Another region distinctly identified by CGH corresponds to an interstitial block localized within a euchromatic chromosome segment. The block was illuminated by both female- and male-derived probes (Fig. 2 j, k, l), which suggests presence of repetitive sequences common to autosomes and Z chromosome (Sahara *et al.* 2003). This block, together with the adjacent terminal region, forms a chromosome segment with distinct molecular composition comprising about one third of the W chromosome. The monarch W thus shows a bipartite organization, with only two-thirds of the chromosome being highly heterochromatic while the remaining third appears euchromatic. Also noteworthy is the large size of the monarch W chromosome relative to chromosomes other than the Z, which is of comparably large size. This combination of size and bipartite organization suggests this chromosome may be a neo-W resulting from a W-autosome fusion occurring in parallel with the neo-Z formation. However, caution is advised in this interpretation because W chromosome size can be misleading (Schartl *et al.* 2016) and cytogenetic examination of species bearing neo-Z chromosomes revealed considerable differences in the structure of their W chromosomes (Šíchová *et al.* 2013).

Finally, it should be noted that a relatively modern W-autosome fusion was recently reported to be segregating in the African Queen butterfly, *D. chrysippus,* where it controls color pattern and male-killing and is driving population divergence across a hybrid zone (Smith *et al.* 2016). Given our results and the same chromosome number N=30 observed in the wild-type *D. chrysippus dorippus,* it seems all *Danaus* species including *D. chrysippus* share the neo-Z and thus presumably this putative neo-W (*i.e.* homologous to McChr21). If so, this means the W-autosome fusion in *D. chrysippus* would be a compound neo-W involving two distinct former autosomes. This pattern of relatively frequent karyotypic changes within the genus further recommends *Danaus* butterflies as an excellent model system for studying sex chromosome evolution.

## Conclusion

We have used a combination of genome sequencing coverage, comparative genomic analysis, and cytogenetics to demonstrate that *Danaus* butterflies harbor a neo-Z chromosome resulting from the fusion of the ancestral Z chromosome and an autosome homologous to Chr21 in *M. cinxia*. Also, at least in the case of Monarch butterflies, it appears that this fusion has resulted in a large neo-W chromosome with a prominent euchromatic region. Our analysis also identified and resolved several Z-autosome chimeric scaffolds in the most recent assembly of the *D. plexippus* genome. This discovery and provisional assignment of chromosomal linkage for >90% of *D. plexippus* genes paves the way for myriad and diverse investigations into sex chromosome evolution, which are likely to be of distinct importance given the increasing prominence of *Danaus* butterflies as a female-heterogametic model species for functional and evolutionary genomics.

## Acknowledgements

Thanks to Chip Taylor, Ann Ryan, and the rest of MonarchWatch.org for donation of organisms used in this study. Jim Mallet and John Davey provided helpful comments on this work. This research was supported by NSF-DEB 1457758 to J.R.W. Cytogenetic analysis was supported by the Czech Science Foundation grant 14-35819P to P.N. The computing for this project was performed on the Community Cluster at the Center for Research Computing at the University of Kansas.

## Authors contributions

AJM, PN, AV, and JRW all performed analyses and contributed to the manuscript text. AJM additionally assembled analyses and manuscript components and edited the complete version. All authors read and approved the final manuscript.

## Additional data files

Monarch_neoZ_Gene-Scaff.xlsx contains a summary of Z-autosome chimeric scaffold breakpoints (Table S3) and the predicted chromosomal linkage, where determined, for all D. plexippus protein coding genes (Table S4).

---

**Figure S1.**
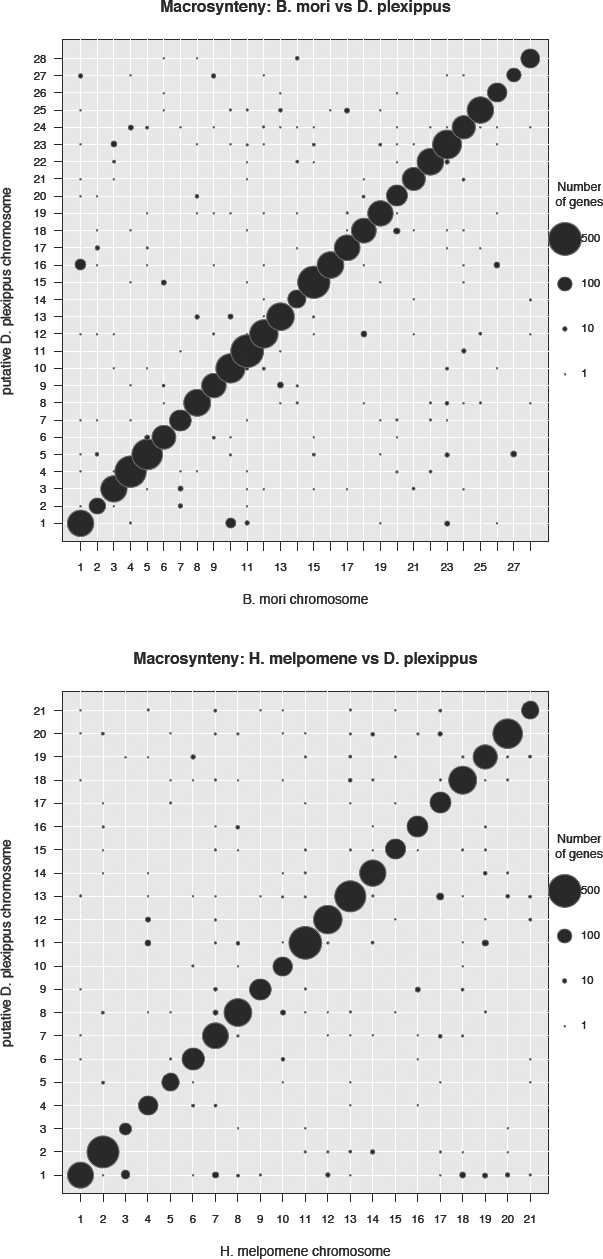
Chromosomal co-linkage between *D. plexippus* and *B. mori* (top) or *H. melpomene* (bottom) for predicted orthologous proteins.

**Figure S2.**
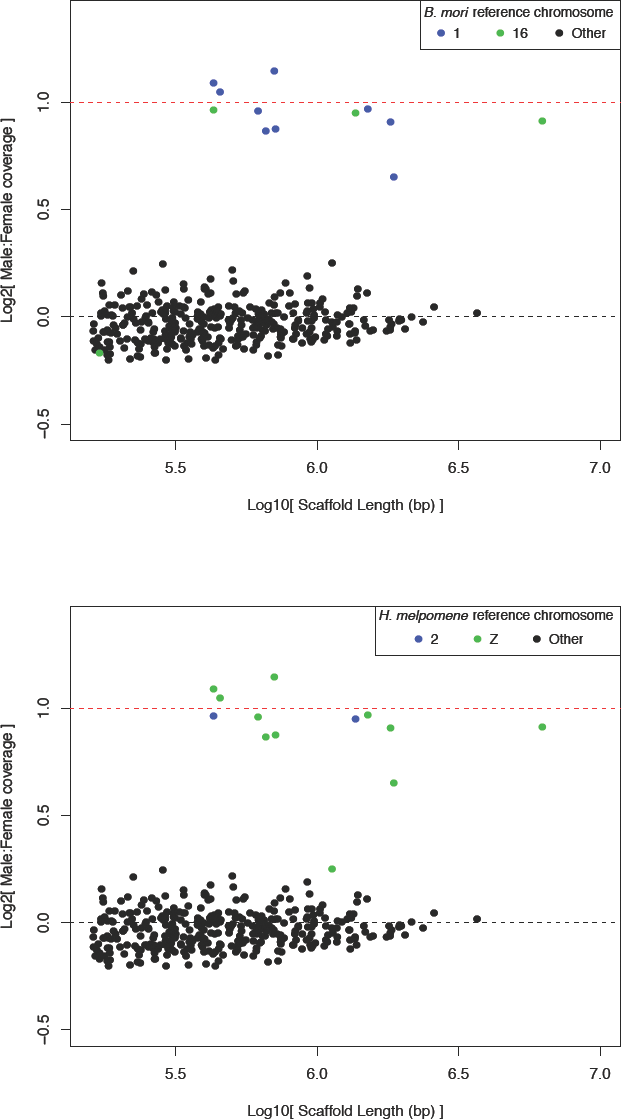
Ratios of male:female median normalized genomic sequencing coverage plotted by scaffold length. Scaffolds assigned to chromosomes putative homologous to the neo-Z chromosome in *D. plexippus* are plotted in distinct colors. Top, relative to *B. mori,* to chromosomes 1 (i.e., Z; blue) and 16 (green). Bottom, relative to *H. melpomene,* chromosomes 1 (*i.e.*, Z; green) and 2 (blue).

**Figure S3.**
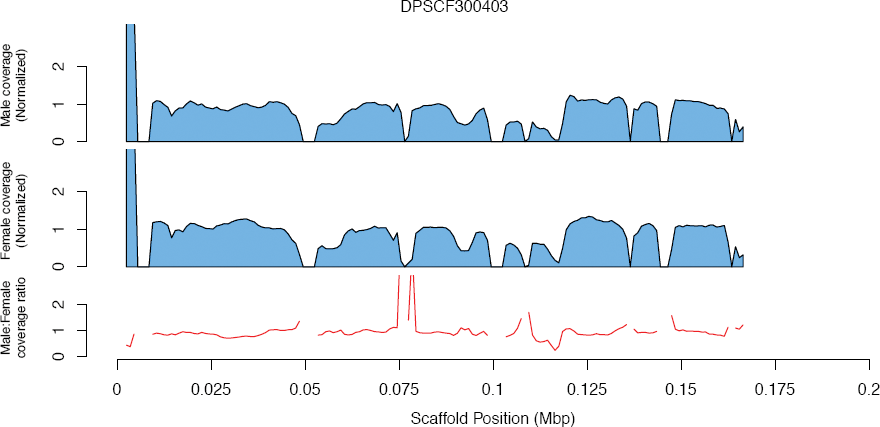
Normalized male and female coverage along the length DPSCF300403. Coverages are plotted as sliding windows (width = 5Kbp, step = 1 Kbp) of median basepair values. The associated male:female ratio of coverage for each window is plotted as a red line below the pair of sex-specific plots.

**Figure S4.**
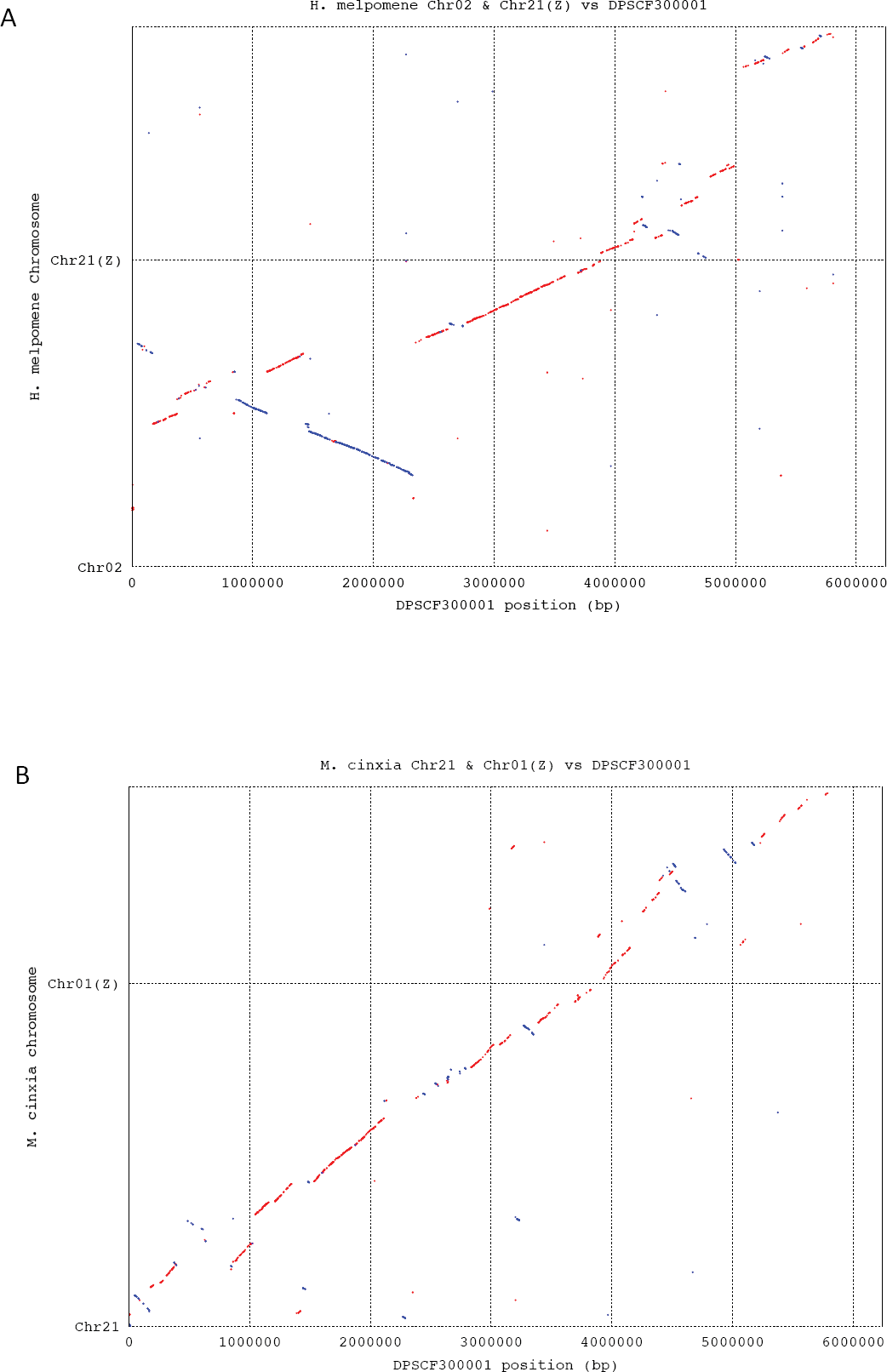
Promer alignments of DPSCF300001 against the Z and homologous autosome from (A) *H. melpomene* and (B) *M. cinxia*. Best one-to-one alignments were generated using default parameters. Manual inspection of alignment coordinates revealed the transition on DPSCF300001 from neo-Z to anc-Z occurs in a window between positions 3.878 and 3.886 Mbp.

**Table S1.**
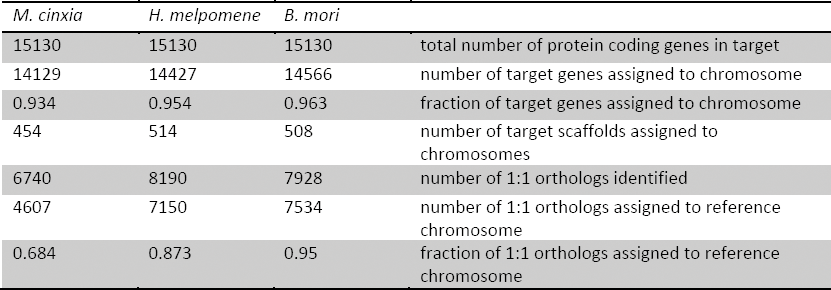
Summary of assigning *D. plexippus* genes and scaffolds to chromosomes via orthology “liftover” relative to three different reference assemblies.

**Supplementary Table S2.** Sample identification details for sequencing data used in coverate analyses.

| Region | Sample | Species | Sex | Collecting location | Date | Accession |
| --- | --- | --- | --- | --- | --- | --- |
| North America | Plex_MA_HI004_M | *plexippus* | male | Massachusetts, USA | July 3 2008 | SRX679269 |
|  | Plex_MA_HI035_F | *plexippus* | female | Massachusetts, USA | August 10 2009 | SRX679310 |
|  | Plex_FLn_StM123_F | *plexippus* | female | St. Marks, Florida, USA | October 2008 | SRX680105 |
|  | Plex_FLn_StM146_M | *plexippus* | male | St. Marks, Florida, USA | October 2009 | SRX681753 |
|  | Plex_WSM_M36_M | *plexippus* | male | Samoa | June 2007 | SRX680118 |
|  | Plex_WSM_M38_F | *plexippus* | female | Samoa | June 2007 | SRX681528 |
| Other *Danaus* | Erip_BRA_16005_F | *erippus* | female | Brazil | September 2010 | SRX682069 |
|  | Erip_BRA_16008_M | *erippus* | male | Brazil | September 2010 | SRX682070 |
|  | Eres_CRC_92_F | *eresimus* | female | Costa Rica | July 24 2010 | SRX682071 |
|  | Eres_FL_27_M | *eresimus* | male | Florida, USA | July 20 2009 | SRX682072 |
|  | Gili_CRC_30_M | *gilippus* | male | Costa Rica | July 24 2010 | SRX682073 |
|  | Gili_TX_01_F | *gilippus* | female | Texas, USA | October 2010 | SRX998564 |

